# Spatiotemporal Propagation of Sensorimotor Beta Bursts Across Adulthood

**DOI:** 10.64898/2026.04.22.720126

**Authors:** Siyu Long, Xiaobo Liu, Georgios D. Mitsis, Marie-Hélène Boudrias

**Affiliations:** Integrated Program in Neuroscience, McGill University, Montréal, Canada; Center for Interdisciplinary Research in Rehabilitation of Greater Montreal (CRIR), Montréal, QC, Canada; Department of Bioengineering, McGill University, Montréal, Canada; Brain Lab, Jewish Rehabilitation Hospital, CISSS-Laval, Laval, Canada; School of Physical and Occupational Therapy, McGill University, Montréal, Canada

## Abstract

Beta activity (13–30 Hz) is a prominent feature of motor control, widely distributed across the cerebral cortex. However, how beta bursts (transient high-amplitude events) propagate along cortical organizational axes, such as the posterior-anterior gradient that coordinates cortical structure and function from sensory to association regions, remains poorly understood. Using magnetoencephalography (MEG) data from the Cambridge Centre for Ageing and Neuroscience (CamCAN) dataset, beta burst propagation was characterized using burst onset timing and optical flow analysis during motor tasks and rest in 573 participants (ages 18-88). Beta bursts propagated systematically along the posterior-anterior cortical axis during motor tasks, with propagation direction reversing between movement phases and exhibiting hemispheric asymmetry. In contrast, the resting state exhibited no consistent spatial organization of beta burst propagation. Propagation patterns in motor tasks significantly correlated with cortical distributions of GABA_A_, cholinergic, and mu-opioid receptors in a hemisphere-specific and phase-dependent manner. Propagation energy was highest in sensorimotor regions and decreased towards more peripheral areas of the cortex. Older adults exhibited significant temporal expansion of beta activity (earlier pre-movement, later post-movement), suggesting a mediation effect of age on reaction time. These results suggest that the propagation of beta bursts is influenced by cortical architecture and may provide a mechanistic explanation for the age-related slowing of motor function.

## Introduction

Beta-band oscillations (13–30 Hz) are a dominant feature of neural activity, detectable across spatial scales ranging from local extracellular recordings to whole-brain magnetoencephalography (MEG) (Engel & Fries, 2010; Shin et al., 2017). These oscillations play a critical role in motor control, with their dynamics closely linked to movement preparation, execution, and cancellation (Lemon, 2008). Characteristic patterns of movement-related beta desynchronization (MRBD) during movement and post-movement beta rebound (PMBR) have been observed across distributed brain regions, including sensorimotor and prefrontal cortices, as well as subcortical structures such as the thalamus, basal ganglia, and cerebellum (Pfurtscheller et al., 2003; Picazio et al., 2014; Sherman et al., 2016; Wagner et al., 2018; Xifra-Porxas et al., 2019). Recent evidence suggests that beta oscillations manifest as transient, high-amplitude bursts rather than sustained rhythmic activity (Feingold et al., 2015; Sherman et al., 2016). Critically, increases in trial-averaged beta spectral power are driven by changes in the occurrence rate of these discrete burst events (Shin et al., 2017). When averaged across trials, beta burst probability closely corresponds to the sustained desynchronization and synchronization patterns observed in conventional analyses (Little et al., 2019). This suggests that trial-averaged burst metrics offer a robust measure of beta dynamics, preserving the functional signatures established in traditional analyses. Specifically, pre-movement burst rates have been shown to reflect motor inhibition and predict movement initiation speed, while post-movement burst patterns are modulated by movement accuracy (Little et al., 2019; Torrecillos et al., 2018). Moreover, averaged beta burst measures capture activity across the same distributed networks identified when using conventional power analyses, spanning sensorimotor, prefrontal, and subcortical structures (Chikermane et al., 2024; Schmidt et al., 2019; Wessel, 2020).

The neural architecture underlying motor control comprises interconnected circuits that link the motor cortex, basal ganglia, prefrontal regions, cerebellum, and other areas (Debaere et al., 2003; Lemon, 2008). Beta oscillations propagate dynamically through these motor networks, serving as a key signalling mechanism for motor state regulation. Evidence shows that beta bursts travel as spatiotemporal waves across the sensorimotor cortex, with propagation patterns reflecting underlying circuit organization and behavioral demands (Rubino et al., 2006; Zich et al., 2023). During motor inhibitory control, beta bursts follow a hierarchical progression through cortico-subcortical pathways, originating in the subthalamic nucleus before reaching the motor thalamus and sensorimotor cortex (Diesburg et al., 2021). Studies combining neural and behavioral measures further reveal that beta bursts exhibit distinct spatiotemporal patterns during different motor states: predominating bilaterally in sensorimotor regions before movement, lateralizing during initiation, and increasing frontocentrally during successful stopping (Wessel, 2020). Together, these findings demonstrate that beta burst propagation is organized across distributed motor networks, varying with both anatomical pathways and task demands.

The posterior-to-anterior axis represents a fundamental organizational principle of cortical function, reflecting a large-scale functional hierarchy from sensory processing to motor execution and cognitive control (Huntenburg et al., 2018; Margulies et al., 2016). This axis is characterized by systematic variations in structural and functional properties, with posterior regions exhibiting high neuronal density and engaging primarily in sensory processing, while anterior regions contain fewer but more extensively connected neurons involved in motor planning and executive control (Cahalane et al., 2012; Charvet & Finlay, 2014; Elston, 2000). This organizational pattern is further supported by graded variations in cortical myelination and microstructural properties along the axis (Paquola et al., 2019), suggesting that the posterior-anterior gradient may facilitate coordinated information flow from sensory to motor regions. However, despite extensive characterization of vertical subcortical-cortical pathways in motor function (Sherman et al., 2016), the organization of cortical propagation along the posterior-anterior axis remains largely unexplored, particularly for motor-related oscillatory dynamics such as beta activity related to motor control.

Cortical oscillation patterns are known to change with advancing age in healthy populations (Rossini et al., 2007). In the motor cortex, healthy aging is associated with elevated baseline beta power and greater movement-related desynchronization, suggesting age-dependent modulation of GABAergic inhibition (Rossiter et al., 2014). Evidence suggests that age-related increases in resting beta activity may be compensated by a more pronounced movement-related beta modulation, with their combined dynamics predicting behavioral outcomes across the lifespan (Heinrichs-Graham et al., 2018). Furthermore, the spatial organization of beta bursts also undergoes age-related changes. Source localization studies demonstrate shifts in the distribution of beta bursts toward frontal cortical areas with advancing age, particularly after 60 years, with burst rate serving as the principal factor underlying age-related changes in movement-evoked beta activity (Brady et al., 2020; Power & Bardouille, 2021). Despite these insights into age-related changes in beta burst characteristics and regional distribution, it remains unknown how aging modulates their propagation patterns across the cortical surface. Characterizing these age-related changes in whole-brain beta burst dynamics is essential for understanding the neural basis of motor aging and identifying potential markers of healthy versus pathological aging.

In this study, we aim to characterize posterior-anterior beta burst propagation patterns across the cerebral cortex and examine how these dynamics are modulated by aging. First, beta burst propagation pathways were tracked by analyzing spatial and temporal onset patterns across cortical regions. Second, the identified propagation patterns were validated by employing optical flow analysis to independently confirm the directionality and spatial organization of beta burst propagation across the cortical surface. Subsequently, the beta burst propagation patterns were correlated with GABA_A_, cholinergic, and mu-opioid receptor density to identify neurochemical correlates of spatial organization. Finally, age-related changes in beta burst onset timing were quantified, and their relationship with motor performance metrics was investigated. By integrating these complementary approaches, the present work represents, to our knowledge, the first attempt to delineate transient beta burst propagation across the cortex, providing insights into inhibitory control networks and their modulation by aging.

## Materials and methods

### Participants & experimental paradigm

We initially included six hundred and eight participants who had MEG data obtained in Phase 2 of the Cam-CAN examination of healthy cognitive ageing (Taylor et al., 2017). Participants who were left-handed or those in whom beta burst activity could not be reliably detected were excluded, resulting in a final sample of 573 right-handed participants aged 18 to 88 years (mean = 55.16, SD = 18.10), comprising 285 males and 288 females. Each participant performed the ‘Sensorimotor task’ and a ‘Resting state’ scan (Shafto et al., 2014; Taylor et al., 2017). In the sensorimotor task, participants responded with a right index finger button press to unimodal or bimodal audio/visual stimuli. The order of bimodal and unimodal trials was randomized, and the inter-trial interval varied between 2 and 26 s.

### Data acquisition

Data were acquired from the Cam-CAN repository (available at http://www.mrc-cbu.cam.ac.uk/datasets/camcan/). MEG recordings were conducted at the MRC Cognition and Brain Sciences Unit (MRC-CBSU, Cambridge) using a 306-channel VectorView MEG system (Elekta Neuromag, Helsinki), comprising 102 magnetometers and 204 orthogonal planar gradiometers housed within a magnetically shielded room (MSR). Data were sampled at 1000 Hz and band-pass filtered online between 0.03 and 330 Hz. Continuous head position tracking within the MEG helmet was enabled by four Head Position Indicator (HPI) coils, allowing for offline motion correction. Vertical and horizontal electrooculogram (VEOG, HEOG) signals were recorded using bipolar electrodes to monitor eye blinks and saccades, while a separate bipolar electrode pair recorded electrocardiogram (ECG) signals to detect pulse-related artifacts. The MEG protocol included approximately 12 minutes of sensorimotor task and 8 minutes and 40 seconds of resting-state data collection. Structural T1-weighted magnetic resonance images (MRI) were obtained using a 3T Siemens Tim Trio scanner equipped with a 32-channel head coil.

### MEG preprocessing

MEG data preprocessing and analysis were conducted using Brainstorm (Tadel et al., 2011) with default settings unless stated otherwise, adhering to established preprocessing guidelines. The preprocessing pipeline consisted of several stages to remove artifacts while preserving neural signals of interest. Raw MEG signals first underwent high-pass filtering using a finite impulse response (FIR) filter at 0.3 Hz to eliminate slow-wave drift and DC offset. Following visual inspection to identify problematic channels and contaminated time segments, power-line interference at 50 Hz and its harmonics were suppressed via notch filtering. A low-pass filter (0.6-280 Hz) was then applied to attenuate high-frequency noise. Physiological artifacts arising from cardiac and ocular activity were removed using signal-space projection (SSP), with artifact components defined based on electrocardiogram (ECG) and electrooculogram (EOG) recordings.

### Trial segmentation and averaging

Task-related MEG data were segmented into trials based on the finger tapping response. Each trial included a 1.4 s pre-movement phase (before button press) and a 1.4 s post-movement phase (after button press, Fig. 1A left). Signals were maintained at 1,000 Hz to preserve precise temporal resolution for accurate beta burst onset time detection. Trials with inter-response intervals less than 3s were excluded to prevent temporal overlap. The remaining trials were averaged across trials for each sensor and participant, producing a single averaged trial per sensor. This approach enhanced the signal-to-noise ratio for source reconstruction and optical flow analysis, enabling reliable detection of propagation direction and energy gradients. While single-trial analysis is valuable for examining trial-by-trial behavioral variability (Little et al., 2019), our focus on spatial organization required robust estimates of propagation patterns that are reproducible across participants, which was enhanced by trial averaging.

**Fig. 1.**
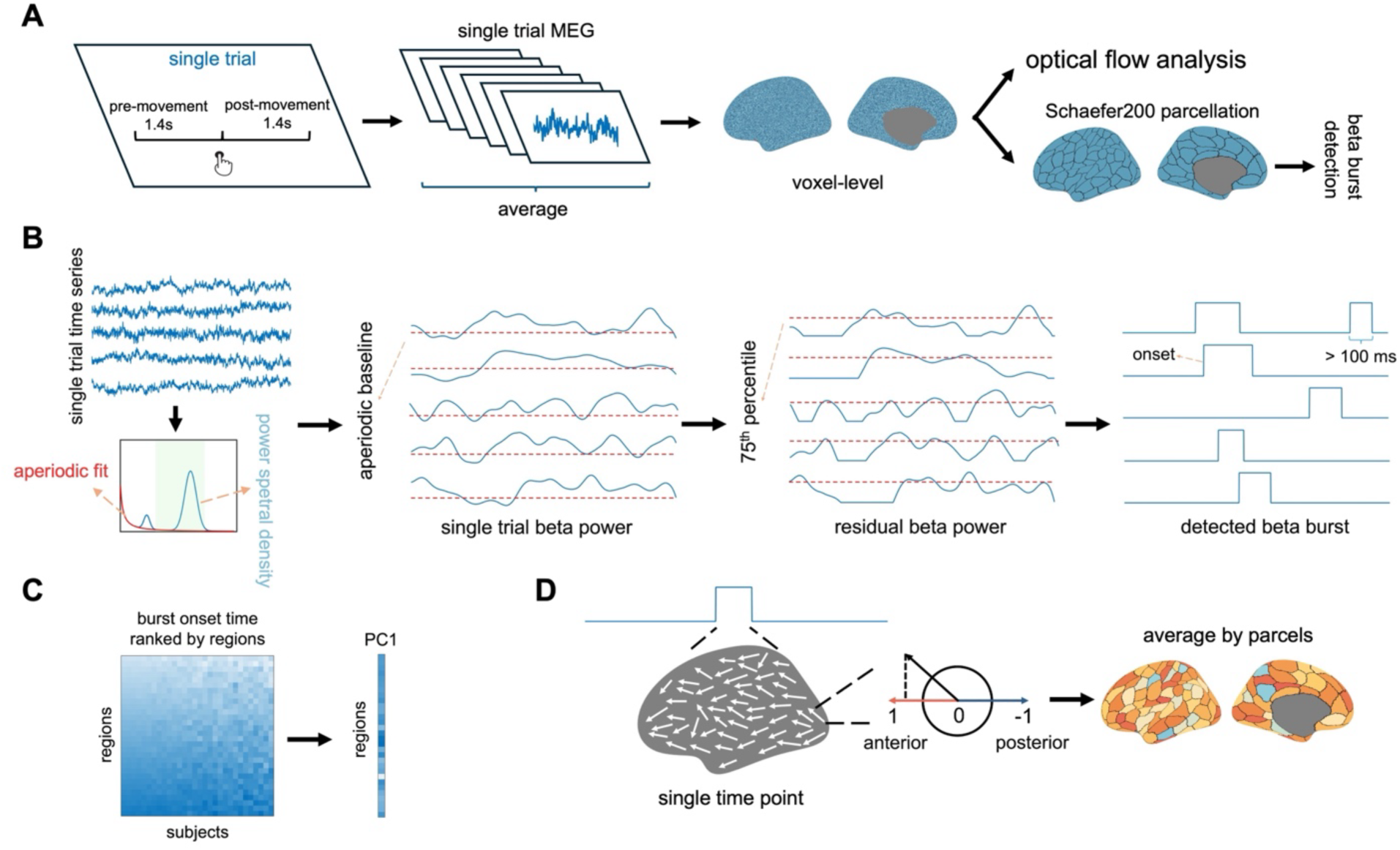
Beta propagation processing pipeline. **(A)** Trial segmentation and source reconstruction workflow. Single trials were defined as the interval between 1.4 s pre-movement and 1.4 s post-movement phases relative to finger tapping. Trial-averaged MEG data were source-reconstructed to 15,000 cortical vertices for optical flow analysis and parcellated to 200 regions for beta burst detection. **(B)** Beta burst detection pipeline. Regional time series underwent time-frequency decomposition, aperiodic baseline subtraction, and amplitude thresholding (75^th^ percentile) to identify bursts exceeding 100 ms duration. **(C)** Burst onset timing analysis. Regional onset times were ranked across participants and subjected to principal component analysis to extract spatial patterns (PC1). **(D)** Optical flow analysis. Flow direction and energy were computed at each cortical vertex and time point, then projected onto the posterior-anterior axis and averaged within parcels.

### Source reconstruction

Source reconstruction was performed using individual T1-weighted MRI scans. Cortical surfaces were reconstructed and tissue-segmented using FreeSurfer (Fischl, 2012), then tessellated into triangular meshes. MEG sensor positions were co-registered to individual anatomy via digitized scalp landmarks. Source activity was estimated using linearly constrained minimum variance (LCMV) beamforming with an overlapping spheres forward model (Veen et al., 1997). Empty-room noise covariance matrices were used to normalize source estimates and correct for depth bias. In total, 15,000 current dipoles were estimated across the cortical surface. The high-resolution source signals were used directly for optical flow analysis to capture fine-grained spatiotemporal dynamics (Fig. 1A – right panel). For beta burst detection and other analyses, source data were parcellated using the Schaefer200 atlas, with vertices within each parcel averaged to create regional time series.

### Beta burst detection

Beta bursts were identified from source-reconstructed time series using a threshold-based detection method (Fig. 1B). For each phase (pre- and post-movement), time-frequency decomposition was performed using the Morlet wavelet transform with a 1 Hz central frequency and a 1-second time resolution. The resulting time-frequency power was averaged across the beta frequency range (13-30 Hz) to obtain a single time-varying beta power signal. To isolate oscillatory activity from aperiodic background activity, we estimated the time-resolved aperiodic component using the SPRiNT (Spectral Parameterization Resolved in Time) algorithm (Wilson et al., 2022). SPRiNT extends the FOOOF (Fitting Oscillations & One-Over-f) approach (Donoghue et al., 2020) to the time domain by computing short-time Fourier transforms (STFT) for sliding time windows, converting the resulting complex frequency domain signals to power by taking the squared magnitude, and performing FOOOF fitting on the log-transformed power spectra. For each phase, we applied SPRiNT with one-second sliding time windows and no overlap to extract the time-varying aperiodic component. The aperiodic fit was then averaged within 13-30 Hz to derive a movement phase-specific aperiodic baseline that accounts for non-oscillatory power fluctuations. For burst detection, the aperiodic baseline was subtracted from the beta time-frequency signal for each movement phase, setting negative values to zero. This subtraction isolates bursts with amplitudes above those of the background neural activity, ensuring that only periods where beta oscillatory power exceeds the aperiodic background are retained (Szul et al., 2023). A ‘flexible interval’ approach was employed for burst detection to avoid edge effects and ensure complete burst identification. An initial time window, ranging from -1.2 to 1.2 s relative to the finger tapping response, was defined for each phase. The residual beta power was then binarized using a 75^th^ percentile threshold computed within each phase. If a burst was detected at the boundary of the time window (either at the beginning or end), the window was extended until the burst ended, ensuring that all detected bursts were fully captured. To identify discrete bursts, we retained only continuous suprathreshold periods exceeding 100 ms in duration, as shorter events likely reflect noise rather than genuine burst activity. The resulting binary time series indicated the presence (1) or absence (0) of beta bursts at each time point.

To characterize the spatiotemporal organization of beta burst propagation, the timing of burst onset was analyzed relative to the finger tapping response. For each trial and cortical region, the beta burst closest in time to the motor response was identified within each phase (pre- and post-movement). The onset time of this selected burst was extracted for each region, yielding a distribution of onset times across cortical regions for each participant (Fig. 1C). To identify common propagation patterns across participants, principal component analysis (PCA) was applied to regional burst onset times. Given the unilateral finger tapping of the motor task, analyses were conducted separately for each hemisphere and movement phase. For each participant, regional onset times were vectorized. PCA was applied to these vectors across participants to extract principal components capturing systematic spatial patterns of burst onset timing. PC1 coefficients for each region represent the relative timing of burst onset, with the spatial distribution of these coefficients characterizing the direction of propagation across cortical regions.

### Optical flow

The optical flow of source-reconstructed beta cortical activity was estimated using a geodesic cortical flow framework (Liu et al., 2026). This approach models the cortical surface as a two-dimensional Riemannian manifold and treats source-imaged MEG activity as a time-varying scalar field defined on this manifold. At each cortical vertex and time point, a surface-tangent propagation vector is estimated by solving an optical flow problem adapted to curved surfaces (Lefèvre & Baillet, 2008; Lefèvre & Baillet, 2009). Specifically, the method assumes local conservation of the scalar field along flow trajectories and estimates the cortical flow vector field by minimizing an energy functional that balances fidelity to the observed spatiotemporal dynamics with a spatial smoothness regularization term. The resulting vector at each vertex encodes both the direction and magnitude of local propagation. To enable consistent interpretation of propagation direction across the folded cortical surface, flow vectors were expressed in an anatomy-informed geodesic reference frame defined on each vertex’s tangent plane, with orthogonal posterior-anterior and superior-inferior axes derived from hemisphere-specific anatomical landmarks on the FreeSurfer FsAverage template surface (Fischl et al., 1999). Flow magnitude was summarized as kinetic energy, defined as the squared norm of the cortical flow vector. This framework has been validated using synthetic simulations with known propagation trajectories and replicated across independent MEG datasets (Liu et al., 2026).

The optical flow algorithm computed flow direction (angle) and flow energy (magnitude) at each cortical voxel and time point. Flow angles were defined such that 0 radians represents posterior-to-anterior flow and π radians represents anterior-to-posterior flow. To quantify propagation along the posterior-anterior axis, we projected the flow angle at each voxel and time point onto this axis (Fig. 1D). These projected angles were then averaged across voxels within each region of the Schaefer200 atlas to obtain region-wise directional estimates.

To focus on the period of active burst propagation, a participant-specific time window was defined based on burst onset timing. For each participant, the earliest burst onset time across all cortical regions was identified, and a 100 ms window starting from this time point was defined. This window length was chosen based on the observed temporal spread of burst onset across regions. The difference between the earliest and latest burst onset times within each participant, when averaged across participants, was approximately 100 ms (Fig. S1), indicating that this window captures the primary propagation period. Optical flow measures were computed within this time window for subsequent analyses.

For the optical flow direction, flow angles were characterized in two ways. First, to visualize the overall spatial pattern of propagation direction, flow angles were averaged across time points within the 100 ms window for each region and participant, yielding spatial distribution maps. Second, to characterize the full distribution of flow directions without temporal averaging, we computed probability density distributions of flow angles across all voxels and time points within the window, providing a comprehensive view of directional variability.

For energy analysis, flow energy was averaged across voxels within each Schaefer200 region and across the 100 ms time window, yielding a single energy value per region for each participant. We then performed PCA on these region-wise energy values to characterize the spatial organization of propagation strength across the cortex.

### Resting state beta burst extraction

To determine whether propagation patterns are task-specific, resting-state beta bursts were analyzed using the same preprocessing and source reconstruction procedures as for task data. Continuous resting-state recordings were segmented into non-overlapping 4-second epochs, and burst detection followed the same pipeline as for task data.

For onset timing analysis, a reference region was first identified as the region with the highest number of bursts across all participants and epochs. For each burst in other regions, the relative onset time was calculated by finding the nearest burst in the reference region and computing the time difference between them. The reference region thus had an onset time of zero by definition. Burst pairs with onset differences exceeding 500 ms (five times the mean task-related onset difference) were excluded to eliminate spurious matches.

### Geodesic distance

To examine the relationship between anatomical location and beta burst propagation patterns, Euclidean distances were computed along two anatomical axes based on region centroid coordinates in the Schaefer200 parcellation. For the posterior-anterior axis, Euclidean distance was computed from the center of the visual cortex to the center of each cortical region (Fig. S2A). For the central-peripheral axis, distance was computed from the center of the motor cortex to each region (Fig. S2B). Only ipsilateral distances were considered, with distances calculated separately for each hemisphere. These distances were then used to examine the spatial distribution of burst onset timing and propagation energy along these axes.

### Receptor map processing

To investigate the potential neurochemical substrates of beta burst propagation patterns, we examined the relationship between regional neurotransmitter receptor densities and the organization of burst onset timing. Three receptor systems were analyzed given their established roles in motor function: γ-Aminobutyric acid (GABA_A_) receptors, which mediate cortical inhibition and are essential for beta oscillation generation in the motor cortex (Nowak et al., 2017; Rossiter et al., 2014); vesicular acetylcholine transporter (VAChT), which is critical for acetylcholine packaging and neuromuscular transmission in motor neurons (Joviano-Santos et al., 2021; Schütz, 2005); and mu-opioid receptors, which provide inhibitory modulation of motor control circuits (Bezard et al., 2020; Wiskerke et al., 2012). Receptor density maps were obtained from publicly available PET datasets (Hansen et al., 2022), consisting of volumetric images aligned to the MNI-ICBM 152 nonlinear 2009 template and parcellated according to the Schaefer200 atlas (https://github.com/netneurolab/hansen_receptors). For each receptor system, we computed spatial correlations between regional receptor density and PC1 coefficients of beta burst onset timing, separately for each hemisphere and movement phase, to assess whether propagation patterns align with cortical distributions of neurotransmitter systems.

### Spatial correlation

To account for spatial autocorrelation in cortical maps, we applied the spatial permutation test (spin test) to all spatial correlation analyses (Markello & Misic, 2021). The spin test generates a null distribution that preserves the spatial autocorrelation of each map while disrupting their correspondence, providing a conservative test of spatial associations. For each correlation (e.g., PC1 vs. spatial distance, PC1 vs. receptor density), cortical surface coordinates were randomly rotated, and regional values were reassigned based on the nearest region after rotation. This procedure was repeated 1,000 times to generate a null distribution. The observed correlation was compared against this distribution to obtain a spatial permutation *p*-value (*p_spin_*). This corrects for the tendency of spatially proximate regions to have similar values, which can inflate correlation coefficients.

### Mediation analysis

To assess whether age-related changes in beta burst onset timing contribute to motor performance, mediation analysis was conducted with age as the independent variable, cortex-averaged onset time as the mediator, and reaction time as the dependent variable. The analysis estimated the direct effect of age on onset time (path a), the effect of onset time on reaction time, controlling for age (path b), and the total (c) and direct (c’) effects of age on reaction time. The indirect effect (ab) quantifies the mediation by onset timing. Complete mediation is indicated when ab is significant and c’ is non-significant (Zhao et al., 2010). Significance was assessed using bootstrap confidence intervals (10,000 iterations). Analyses were performed separately for the pre-movement and post-movement phases.

## Results

### Beta bursts propagate along the posterior-anterior axis with phase-dependent reversal and hemispheric asymmetry

Given that our task involved unimanual right-hand movements, which introduces hemispheric asymmetry between the dominant (left) and non-dominant (right) hemispheres, we performed separate PCA analyses for each hemisphere and movement phase. PCA of regional beta burst onset times revealed systematic spatial organization along anatomical gradients (Fig. 2A-B), with the first principal component (PC1) explaining more than 24% of the variance in both movement phases (Fig. S3). During the pre-movement phase (Fig. 2A, upper panel), the left hemisphere showed negative correlations between PC1 and posterior-to-anterior distance (Fig. 2B, first panel, *ΔAdj. R^2^* = 0.18, *p* = 8.5 × 10^-6^, *p_spin_* = 0.02), while the right hemisphere showed positive correlations (Fig. 2B, second panel, *ΔAdj. R^2^* = 0.46, *p* = 4.3 × 10^-15^, *p_spin_* = 0.01). During the post-movement phase (Fig. 2A, lower panel), this hemispheric asymmetry was reversed, with positive correlations in the left hemisphere (Fig. 2B, third panel, *ΔAdj. R^2^* = 0.79, *p* < 1 × 10^-20^, *p_spin_* < 0.001) and negative correlations in the right hemisphere (Fig. 2B, fourth panel, *ΔAdj. R^2^* = 0.75, *p* < 1 × 10^-20^, *p_spin_* < 0.001). These findings reveal that beta burst propagation follows organized spatial gradients along the posterior-to-anterior axis, with opposite hemisphere-specific directional patterns between movement phases.

**Fig. 2.**
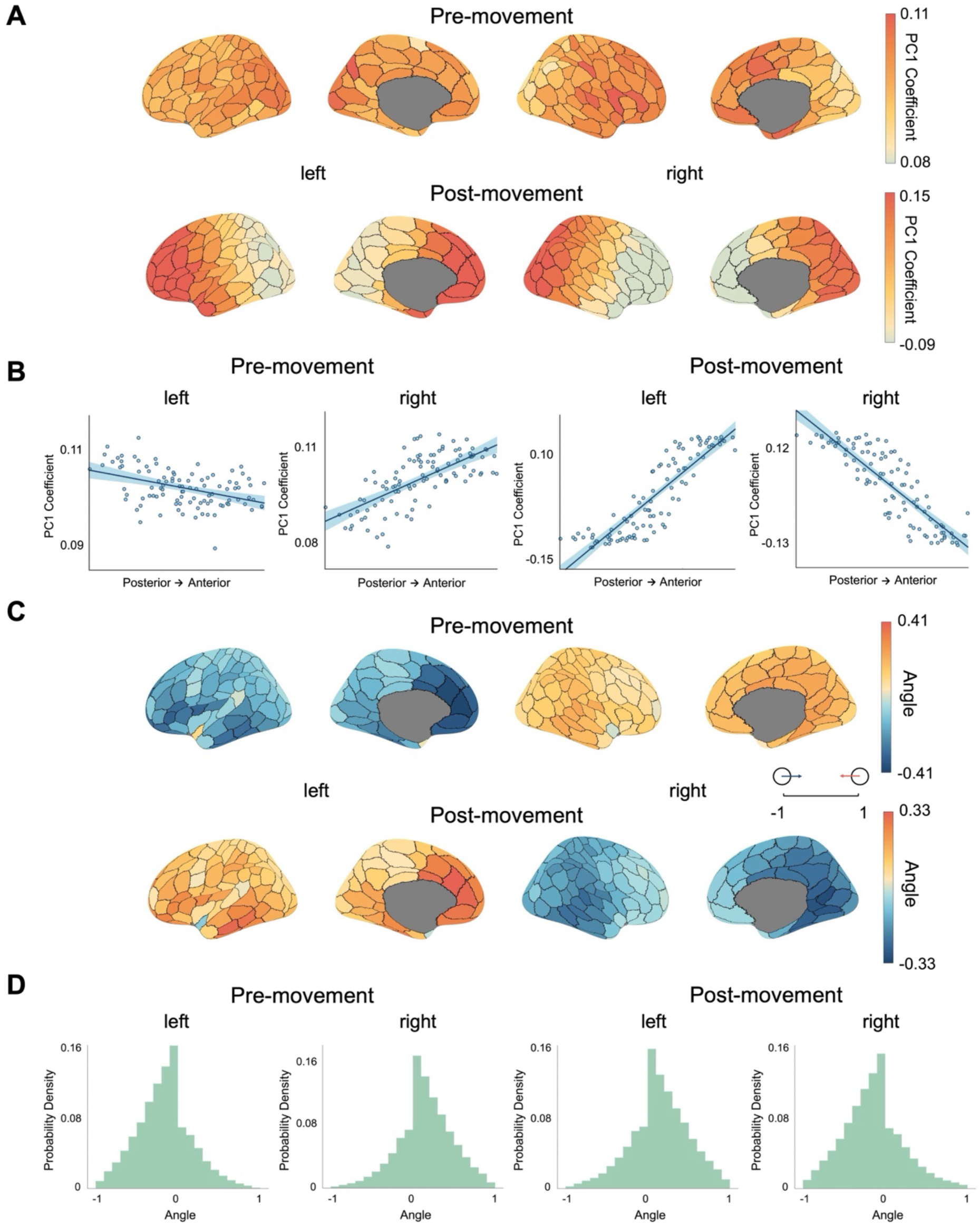
Spatial organization of beta burst propagation along the posterior-anterior axis. **(A)** Cortical maps showing PC1 of regional beta burst onset times during pre-movement (top) and post-movement (bottom) phases. **(B)** Scatterplots displaying correlations between PC1 coefficients and posterior-to-anterior distance for each hemisphere (left and right panels) during pre-movement (left two plots) and post-movement (right two plots). Beta burst onset times were significantly correlated with posterior-to-anterior distance (all *p_spin_* < 0.05), with hemisphere-specific patterns that reversed between movement phases. **(C)** Cortical maps showing participant-averaged optical flow directions projected onto the anterior-posterior axis within 100 ms following beta burst onset for pre-movement (top) and post-movement (bottom). Positive values (warm colors) indicate posterior-to-anterior flow; negative values (cool colors) indicate anterior-to-posterior flow. **(D)** Probability density distributions of optical flow angles across all time points, parcels and participants without averaging, for pre-movement (left) and post-movement (right) in each hemisphere. Optical flow patterns validate the spatial organization observed in onset timing analysis.

To provide independent validation of these spatial propagation patterns, optical flow analysis was performed on beta time-frequency dynamics (Fig. 2C-D). During the pre-movement phase (Fig. 2C, upper panel), the spatial maps illustrate optical flow directions averaged across the 100 ms window and all participants, showing hemisphere-specific patterns that corresponded to the PC1 gradients observed in the onset timing analysis. Probability density distributions (Fig. 2D, left panel) show the distribution of optical flow angles across all time points within the 100 ms window without temporal or across-participant averaging. One-sample t-tests confirmed that these distributions exhibited significant directional biases (both *p* < 1 × 10^-20^, one-sample t-tests), with predominantly positive angles in the left hemisphere and negative angles in the right hemisphere during pre-movement. During the post-movement phase (Fig. 2C, lower panel), both the spatial maps and probability density distributions revealed opposite optical flow directions. The corresponding probability density distributions (Fig. 2D, right panel) demonstrated this reversal statistically, with predominantly negative angles in the left hemisphere and positive angles in the right hemisphere (both *p* < 1 × 10^-20^, one-sample t-tests). These optical flow patterns are consistent with the spatial gradients observed for beta burst onset timing, providing convergent evidence for organized posterior-anterior propagation of beta burst activity.

### Beta burst propagation energy decreases from motor cortices to peripheral regions

Beyond propagation direction, the spatial distribution of beta burst propagation velocity was examined to determine whether propagation speed varied systematically across the cortex (Fig. 3). Optical flow energy quantifies the magnitude of propagating activity, reflecting the velocity of the optical flow dynamics. During the pre-movement phase (Fig. 3A, upper), the PC1 of optical flow energy explained 45.68% of the variance and showed a clear spatial organization (Fig. S4, left), with higher energy in motor-related cortical regions (central) and lower energy in peripheral regions. PC1 coefficients negatively correlated with central-to-peripheral distance across both hemispheres (Fig. 3B, left, *ΔAdj. R²* = 0.47, *p* < 1 × 10^-20^, *p_spin_* < 0.001). During the post-movement phase (Fig. 3A, lower), PC1 explained 21.15% of the variance and exhibited a similar spatial organization (Fig. S4, right), with a significant negative correlation between PC1 coefficients and spatial distance (Fig. 3B, right, *ΔAdj. R²* = 0.49, *p* < 1 × 10^-20^, *p_spin_* < 0.001). These findings demonstrate that beta burst propagation velocity follows organized spatial gradients, with stronger propagation in central cortical regions and weaker propagation in peripheral regions, consistent across both movement phases.

**Fig. 3.**
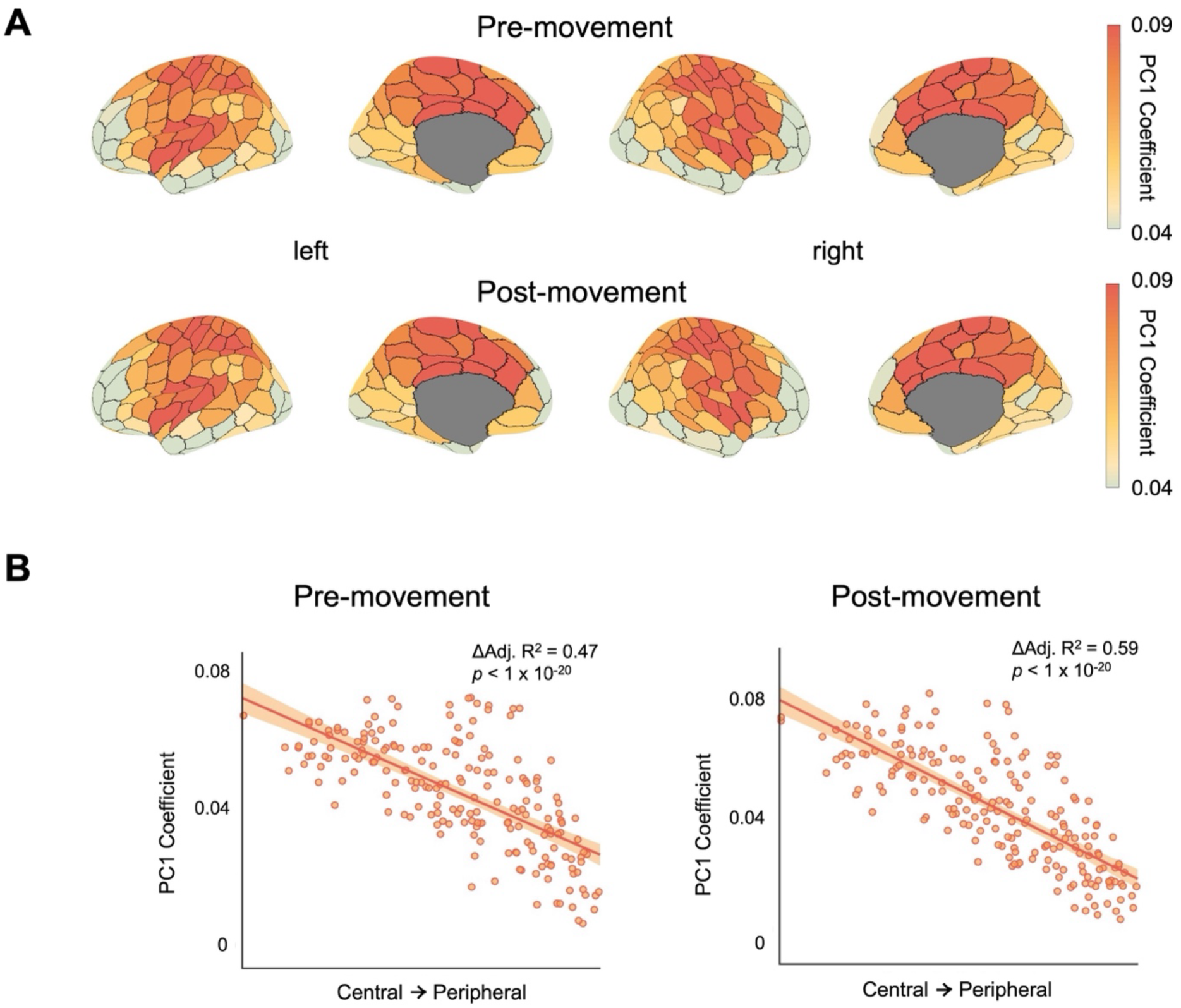
Optical flow energy exhibits a systematic decrease from motor cortices to peripheral regions. **(A)** Cortical surface maps showing the PC1 of optical flow energy for pre-movement (top) and post-movement (bottom) phases across both hemispheres. **(B)** Scatterplots displaying correlations between PC1 coefficients and central-to-peripheral distance pooled across hemispheres for pre-movement (left, *ΔAdj. R²* = 0.47, *p* < 1 × 10^-20^, *p_spin_* < 0.001) and post-movement (right, *ΔAdj. R²* = 0.59, *p* < 1 × 10^-20^, *p_spin_* < 0.001) phases.

### Beta bursts occur randomly during rest

To determine whether the propagation patterns were unique to motor task execution, beta burst onset timing was also examined during the resting state. PCA was performed on regional onset ranks and the PC1 spatial pattern was extracted (Fig. 4A). Unlike the organized patterns observed during the motor task, the resting-state PC1 showed no clear systematic spatial organization. To confirm whether this reflected genuine random spatial variation rather than a weak but consistent pattern, we performed permutation testing. For each participant, we correlated their regional onset rank with the group PC1. To test whether the group-mean correlation significantly exceeded chance, a null distribution was generated using 1,000 permutations. In each permutation, the region-to-onset-rank mapping was randomly shuffled for each participant, and individual correlations with the group PC1 were recalculated and averaged across participants to obtain one permutation group-mean value. During rest (Fig. 4B), the observed correlations did not significantly exceed the permutation null distributions (*p* > 0.9 for both hemispheres), confirming that resting-state patterns were random across participants.

**Fig. 4.**
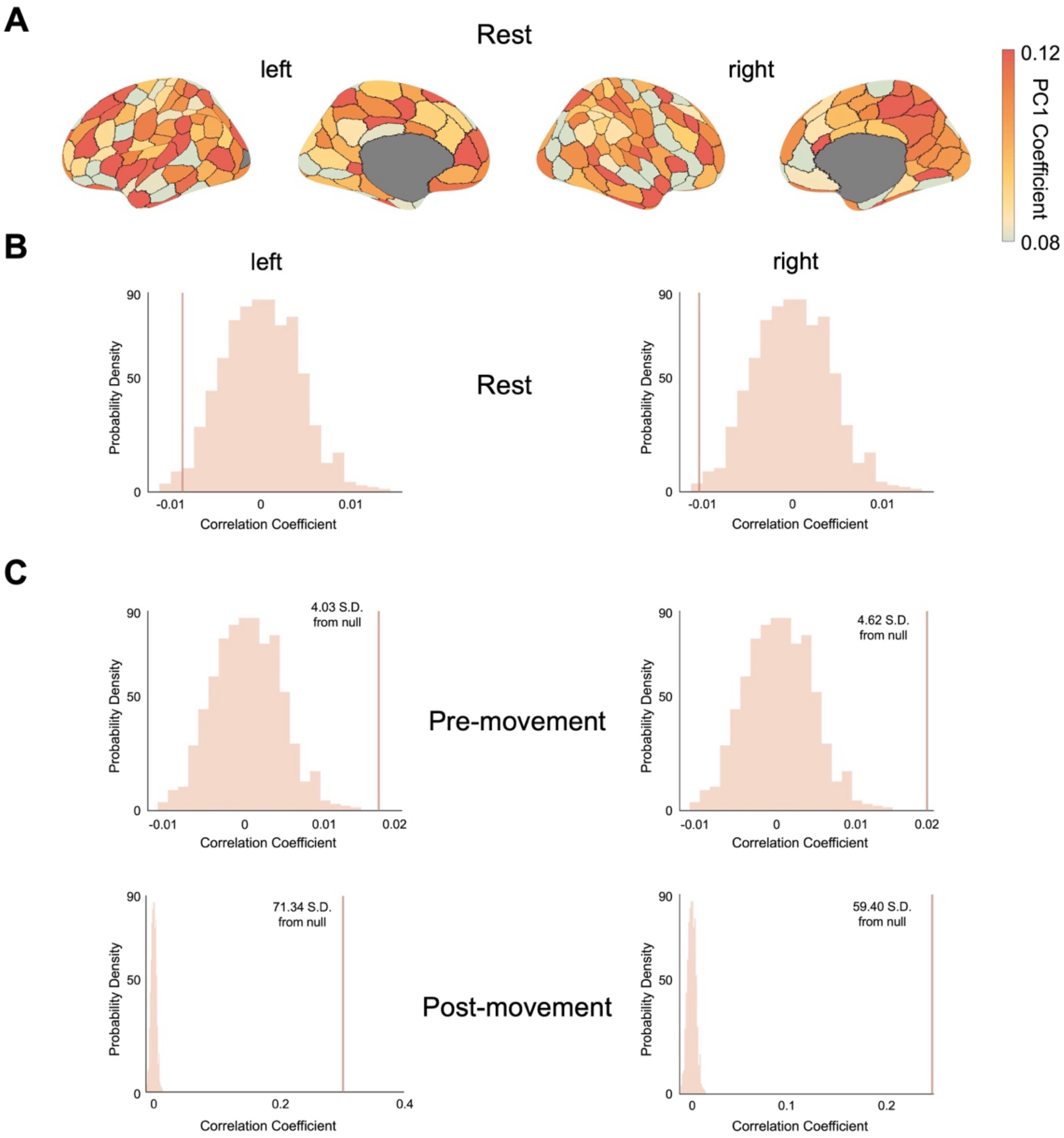
Individual beta burst patterns align with group-level spatial organization during motor task but not rest. **(A)** Group PC1 of regional beta burst onset timing during resting state. **(B)** Permutation null distributions for rest (1,000 iterations). In each iteration, region-onset rank mappings were randomly shuffled for each participant, individual correlations with group PC1 were computed, and then averaged across participants. Vertical lines show the observed mean correlations for the left (left panels) and right (right panels) hemispheres. **(C)** Permutation null distributions for motor task separated by hemisphere (columns) and phase (rows: pre-movement top, post-movement bottom). The observed correlations significantly exceeded permutation distributions in all motor conditions (all *p* < 0.001) but not rest.

The same permutation testing was then applied to the motor task data to contrast with the resting state. During the motor task (Fig. 4C), the observed correlations significantly exceeded chance levels in all conditions (pre-movement left: *z* = 4.03, *p* < 0.001; right: *z* = 4.62, *p* < 0.001; post-movement left: *z* = 71.34, *p* < 0.001; right: *z* = 59.40, *p* < 0.001). This contrast between rest and task demonstrates that organized posterior-anterior propagation patterns emerge specifically during motor execution.

### Beta burst propagation patterns correlate with receptor density

To explore neurochemical substrates of the observed propagation patterns, we examined whether the beta burst onset timing PC1 correlated with regional neurotransmitter receptor densities from published PET atlases (Fig. 5). The associations with GABA_A_, VAChT, and mu-opioid receptor densities were tested across hemispheres and movement phases.

**Fig. 5.**
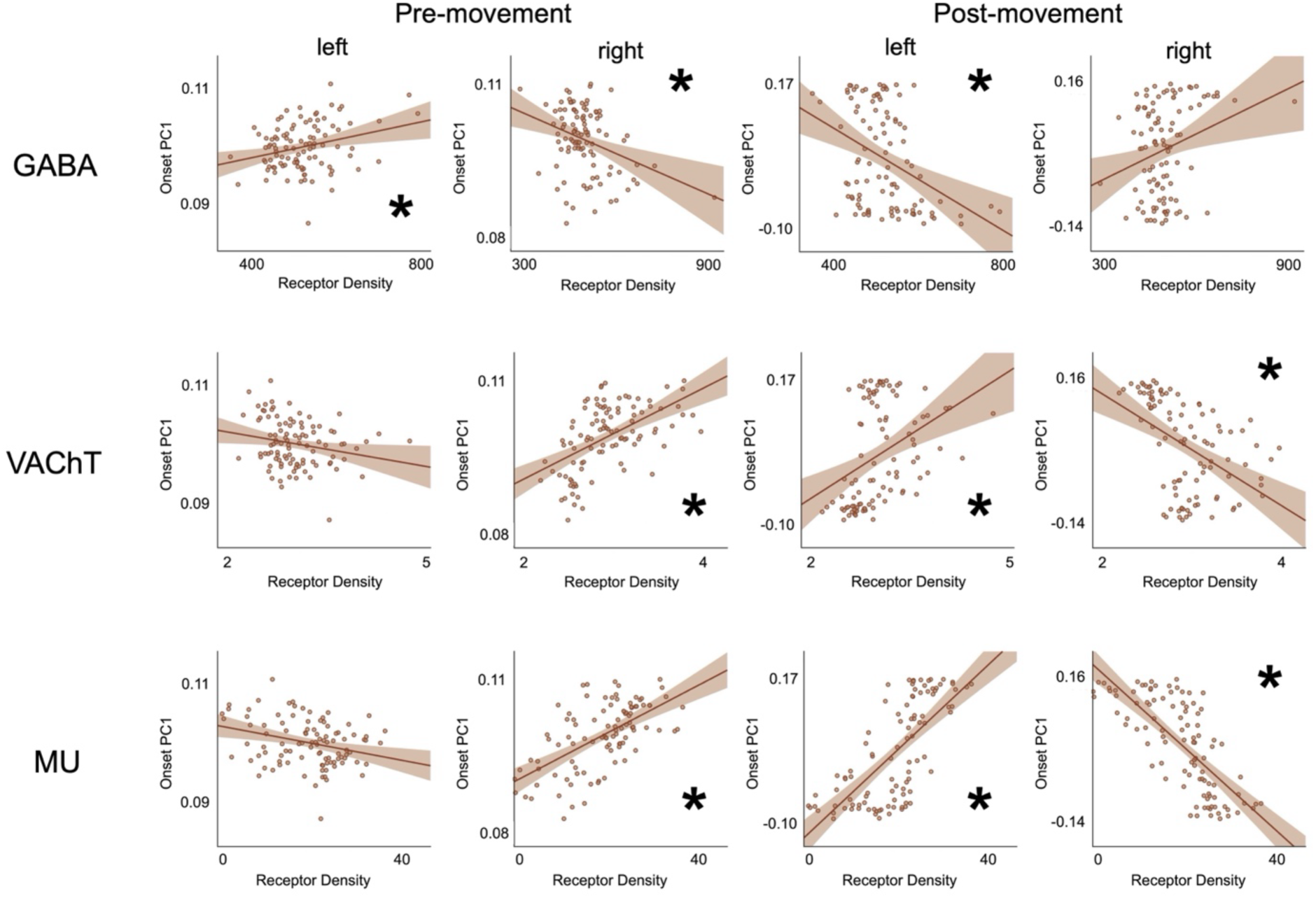
Regional receptor densities correlate with beta burst propagation patterns. PC1 coefficients of onset timing plotted against GABA_A_ (top), VAChT (middle), and mu-opioid (bottom) receptor densities for both hemispheres during pre-movement (left two columns) and post-movement (right two columns). Significant correlations are marked with *.

All three receptor systems showed significant correlations with propagation patterns, exhibiting hemisphere-specific and phase-dependent reversal patterns that paralleled the asymmetric beta burst dynamics. For GABA_A_ receptors, correlations were positive in the left hemisphere during pre-movement (*ΔAdj. R²* = 0.07, *p_spin_* = 0.05) but reversed to negative during post-movement (*ΔAdj. R²* = 0.11, *p_spin_* = 0.02). The right hemisphere showed the opposite pattern, with negative correlations during pre-movement (*ΔAdj. R²* = 0.14, *p_spin_* = 0.04) and positive correlations during post-movement (*ΔAdj. R²* = 0.06, *p_spin_* > 0.05).

VAChT and mu-opioid receptors exhibited similar but complementary patterns. Both showed negative correlations in the left hemisphere during pre-movement (VAChT: *ΔAdj. R²* = 0.04, *p_spin_* > 0.05; mu-opioid: *ΔAdj. R²* = 0.08, *p_spin_* > 0.05) and positive correlations during post-movement (VAChT: *ΔAdj. R²* = 0.14, *p_spin_* = 0.04; mu-opioid: *ΔAdj. R²* = 0.51, *p_spin_* < 0.001). In the right hemisphere, both showed positive correlations during pre-movement (VAChT: *ΔAdj. R²* = 0.29, *p_spin_* < 0.001; mu-opioid: *ΔAdj. R²* = 0.36, *p_spin_* < 0.001) and negative correlations during post-movement (VAChT: *ΔAdj. R²* = 0.22, *p_spin_* = 0.03; mu-opioid: *ΔAdj. R²* = 0.54, *p_spin_* < 0.001). These findings demonstrate that the asymmetric spatial organization of beta burst propagation is closely linked to cortical distributions of multiple neurotransmitter systems.

### Age extends the temporal window of beta bursts

Given that the studied cohort spans a wide age range (18-88 years), we examined whether the posterior-anterior propagation pattern of beta bursts was associated with age. The strength of individual participants’ alignment with the group-level spatial propagation pattern (PC1) showed no significant age-related trend across participants (Fig. S5). We then examined whether age influences the overall timing of beta burst activity by analyzing cortex-averaged onset time across participants (Fig. 6A). Sex was regressed out from onset time prior to analysis. During the pre-movement phase, onset time decreased with age (*Δadj. R²* = 0.02, *p* = 1.3 × 10^-3^), while during the post-movement phase, onset time increased with age (*Δadj. R²* = 0.03, *p* = 3.3 × 10^-5^). These convergent age effects suggest a temporal expansion of beta burst activity around the movement response, with older adults showing beta bursts occurring farther from the movement execution.

**Fig. 6.**
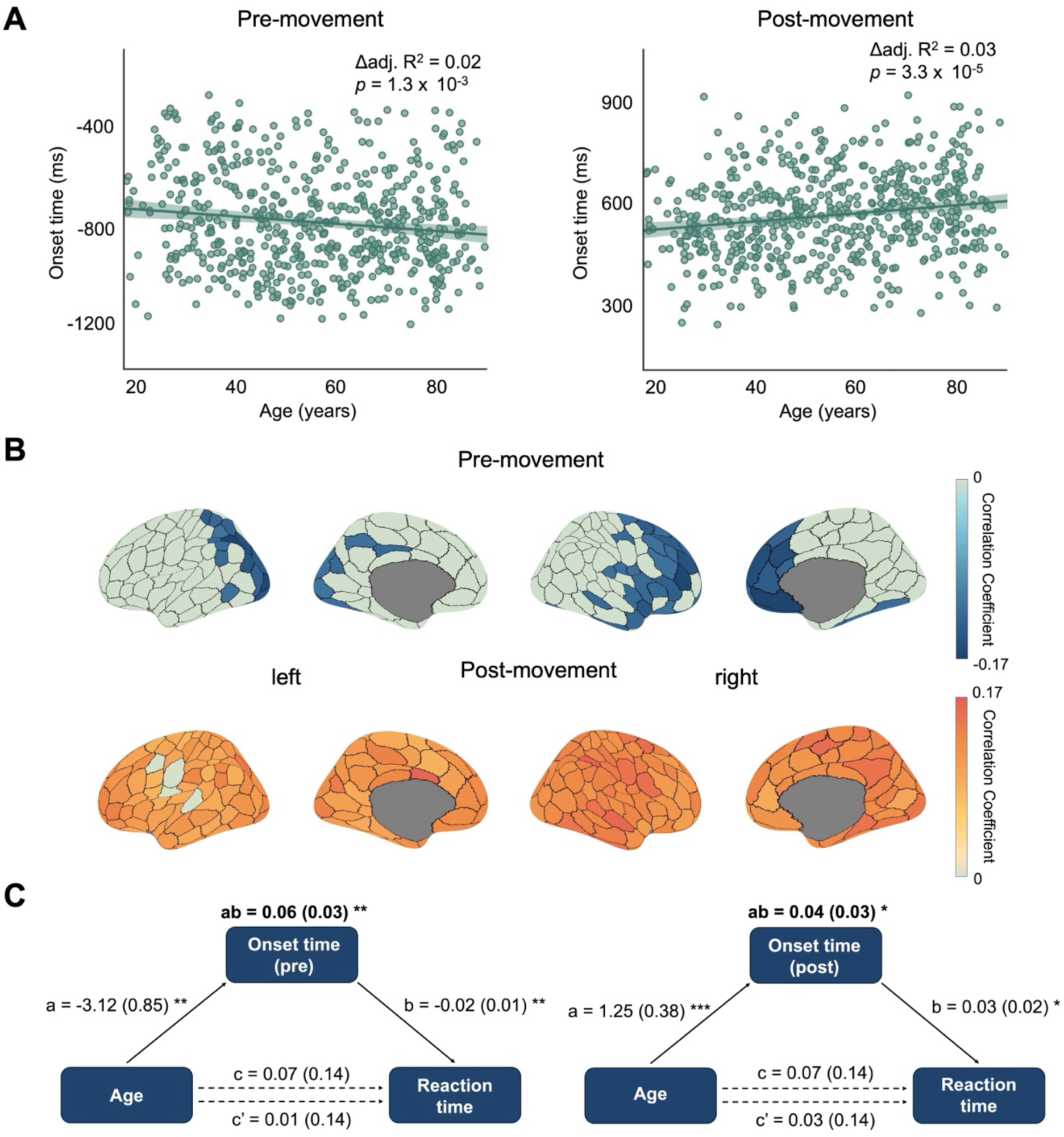
Age-related temporal expansion of beta burst onset timing mediates motor performance changes. **(A)** Scatterplots displaying correlations between cortex-averaged beta burst onset time and age during pre-movement (left, *ΔAdj. R²* = 0.02, *p* = 1.3 × 10^-3^) and post-movement (right, *ΔAdj. R²* = 0.03, *p* = 3.3 × 10^-5^) phases, with sex regressed out as a covariate. Beta onset time showed negative correlations with age during pre-movement and positive correlations during post-movement. **(B)** Cortical maps showing regional correlations between onset time and age. All cortical regions demonstrated consistent correlation directions for both movement phases. Colored regions survived FDR correction; uncolored areas showed similar trends without reaching significance. **(C)** Path analysis demonstrating complete mediation of age effects on reaction time through onset timing changes. Pre-movement (left) and post-movement (right) phases both showed significant indirect effects (ab = 0.06 and 0.04, respectively). Standardized coefficients with standard errors in parentheses; solid lines indicate significant paths.

The spatial distribution of age-related changes was next examined by analyzing regional correlations between onset time and age across the cortex (Fig. 6B). All cortical regions showed consistent correlation directions, with negative correlations during the pre-movement period and positive correlations during the post-movement period, paralleling the whole-brain pattern. Regions surviving False Discovery Rate (FDR) correction were predominantly distributed across frontal and sensorimotor cortices in both phases, while non-significant regions exhibited similar correlation trends with insignificant correlation. This widespread pattern suggests that age-related temporal shifts in beta burst activity represent a global cortical phenomenon.

To determine whether age-related changes in beta burst onset timing contribute to motor performance, we conducted mediation analyses examining the relationship between age, onset time, and reaction time (Fig. 6C). Both movement phases revealed complete mediation. During the pre-movement phase, older age was associated with earlier beta burst onset (a = -3.12, *p* < 0.01), and earlier onset was associated with slower reaction times (b = -0.02, *p* < 0.01). The indirect effect was significant (ab = 0.06, *p* < 0.01), while the direct effect of age on reaction time became non-significant when controlling for onset time (c’ = 0.01, *p* > 0.05). During the post-movement phase, older age was associated with later onset (a = 1.25, *p* < 0.001), and later onset was associated with slower reaction times (b = 0.03, *p* < 0.05), with a significant indirect effect (ab = 0.04, *p* < 0.05) and non-significant direct effect (c’ = 0.03, *p* > 0.05). These complete mediation patterns reveal that prolonged reaction times in older adults are fully accounted for by the temporal expansion of beta burst onset.

## Discussion

In this study, the spatiotemporal organization of beta burst propagation associated with a motor task (right finger button pressing) was characterized in detail. Beta bursts were found to propagate systematically along the posterior-anterior axis during the motor task but not during rest, indicating task-specific spatial organization. This organized propagation reversed direction between the two movement phases (pre and post) and exhibited hemispheric asymmetry, with opposite patterns in contralateral versus ipsilateral hemispheres. The spatial organization of propagation correlated with cortical distributions of GABA_A_, cholinergic (VAChT), and mu-opioid receptors in a hemisphere-specific and movement phase-dependent manner, suggesting neurochemical substrates of the observed propagation patterns. Beyond directional organization, propagation energy was highest in sensorimotor regions and decreased toward the peripheral cortex, reflecting the hub-like role of motor areas. Age-related analysis revealed temporal expansion of beta activity in older participants, with this expansion mediating the relationship between age and reaction time.

### Beta bursts propagation along the posterior-anterior axis

The structural properties of the cortex vary systematically along the posterior-anterior axis, with gradients in neuronal density, connectivity, and myelination (Cahalane et al., 2012; Paquola et al., 2019). These structural gradients are paralleled by functional organization, with the posterior-anterior axis also reflecting hierarchical patterns of cortical processing (Huntenburg et al., 2018; Margulies et al., 2016). This hierarchy characterizes both unimodal sensory processing in posterior regions and transmodal association functions in anterior regions (Margulies et al., 2016). This functional gradient has been also linked to the temporal dynamics of neural activity, whereby the dominant frequency of neural oscillations has been shown to decrease systematically along the posterior-anterior axis, from higher-frequency dynamics in sensory regions to lower-frequency dynamics in association areas (Mahjoory et al., 2020). The posterior-anterior axis has been further associated with directional information flow, with beta oscillations showing posterior-to-anterior propagation from the visual cortex and the posterior default mode network (Hillebrand et al., 2016). Here, beta burst activity was found to propagate systematically along the posterior-anterior axis during motor task performance, demonstrating that this fundamental organizational principle extends to the spatiotemporal dynamics of motor-related neural activity.

Beta oscillations exhibit functionally distinct patterns across movement phases. The MRBD during preparation and execution reflects neural activation and motor engagement, while the PMBR is associated with active motor inhibition, sensory feedback of sensorimotor cortices, and motor state stabilization (Adam et al., 2015; Heinrichs-Graham et al., 2017; Xifra-Porxas et al., 2019). At the level of transient dynamics, beta bursts also show corresponding phase-specific modulations in occurrence rate and functional correlates (Little et al., 2019; Torrecillos et al., 2018). We found that beta burst propagation direction also reversed completely between the pre-movement and post-movement phases, suggesting that the posterior-anterior axis supports bidirectional information flow adapted to these distinct functional demands. This reversal may reflect a shift from feedforward motor processing to feedback-driven sensory integration and motor inhibition, indicating that cortical information flow is dynamically organized rather than following a fixed hierarchical pathway.

Unimanual motor control exhibits pronounced hemispheric asymmetry. The hemisphere contralateral to the moving hand shows stronger beta desynchronization during movement than the ipsilateral hemisphere, with post-movement beta rebound often confined to the contralateral side relative to the moving hand (van Wijk et al., 2012). Beyond amplitude differences, the two hemispheres serve distinct functional roles: beta desynchronization in the hemisphere contralateral to the moving hand reflects neural activation underlying movement execution, whereas ipsilateral beta modulation is associated with motor inhibition and the suppression of unintended movements in the non-task hand (Hosaka et al., 2015). This asymmetry extends to frequency-specific lateralization, with gamma oscillations linked to contralateral motor control and beta oscillations related to ipsilateral motor suppression (Gross et al., 2005; Hosaka et al., 2015). Critically, we found that hemispheric asymmetry extends beyond amplitude and frequency to the spatial organization of propagation: the left and right hemispheres exhibited opposite beta burst propagation directions during both movement phases. This opposite organization pattern demonstrates that the hemispheres are not simply mirror images with different activation strengths, but employ fundamentally different spatial-temporal processing strategies, with each hemisphere organizing information flow in opposite directions along the posterior-anterior axis.

### Propagation energy decreased from motor cortex to peripheral regions

Beta activity propagates continually across the cortex at a range of speeds (Rubino et al., 2006). Motor-related cortical areas are extensively connected with distributed brain networks including sensorimotor, attention, and cognitive control systems (Alahmadi, 2024), and exhibit directional interactions with these distributed regions through beta oscillatory dynamics during movement (Brovelli et al., 2004). These connectivity patterns have established the motor cortex as a coordinating hub within sensorimotor networks. We found that beta burst propagation energy reflected this hub organization, showing highest values in sensorimotor regions with a progressive decrease toward peripheral cortex. Importantly, this center-periphery energy gradient was independent of propagation direction along the posterior-anterior axis. While directional flow varied with movement phase and hemisphere, propagation strength consistently diminished radially from the motor cortex. This spatial organization suggests that the motor cortex functions as a central coordinator of beta burst activity, with the energy gradient indexing the spatial extent of its influence over distributed cortical networks.

### Systematic propagation occurred during motor tasks but not during rest

During the resting state, beta bursts occur through transient co-activations across distributed regions without following a fixed temporal sequence, reflecting spontaneous network dynamics rather than organized propagation (Fries, 2005; Lundqvist et al., 2024). This temporal irregularity is a fundamental characteristic of resting-state beta dynamics, in which beta bursts occur in a largely stochastic manner without being time-locked to specific events. In contrast, during motor preparation and execution, beta bursts exhibit more structured temporal patterns, with their occurrence tightly linked to task-related events (West et al., 2023). Task engagement imposes organizational constraints that transform spontaneous, variable activity into coordinated patterns aligned with behavioral demands. Consistent with this principle, we found that spatial propagation organization also differs between rest and task. Resting-state beta bursts showed no consistent propagation patterns across participants, with burst onset appearing randomly distributed across cortical regions. Motor task engagement, however, produced highly consistent propagation along the posterior-anterior axis with systematic spatiotemporal sequences replicated across trials and participants. This rest-to-task transition demonstrates that systematic propagation is not a spontaneous property of beta bursts but rather emerges in response to the functional demands of motor control, requiring coordinated information flow along cortical organizational axes.

### Neurochemical correlates of beta burst propagation

GABA_A_ receptors play a critical role in motor cortical beta dynamics, mediating spontaneous beta oscillations and modulating movement-related beta changes (Gaetz et al., 2011; Hall et al., 2011). Post-movement beta rebound, in particular, is associated with GABA_A_-mediated inhibition, with stronger rebound reflecting increased inhibitory activity (Zhang et al., 2024). Moreover, propagating beta waves across the motor cortex are controlled by local GABA_A_ inhibition (Rubino et al., 2006), indicating that GABAergic activity regulates not only the temporal dynamics but also the spatial propagation of beta oscillations. Beyond GABA_A_, other neurotransmitter systems also modulate motor function. VAChT is associated with cholinergic innervation of motor neurons (Nagao et al., 1998; Schütz, 2005), while mu-opioid receptors provide inhibitory modulation of motor circuits and ameliorate Parkinsonian motor symptoms (Bezard et al., 2020; Wiskerke et al., 2012). In our study, all three receptor systems (GABA_A_, VAChT, and mu-opioid) showed significant correlations with beta burst propagation patterns in a hemisphere-specific and phase-dependent manner. Notably, the direction of these correlations reversed between movement phases and differed between hemispheres, forming complementary patterns across conditions. This convergence of multiple neurotransmitter systems suggests that beta burst propagation patterns are constrained by the underlying neurochemical architecture. The hemisphere-specific and movement phase-dependent reversal of correlation patterns may reflect the distinct functional roles of each hemisphere and movement phase, with different neurotransmitter balances supporting motor execution versus motor termination, and contralateral versus ipsilateral processing.

### Aging effects on beta burst onset timing

The functional organization of beta burst activity changes across the lifespan. Age-related changes in beta burst dynamics include alterations in burst rate during motor tasks (Brady et al., 2020), shifts in spatial localization with prominent effects in the frontal cortex (Power & Bardouille, 2021), and refinement of temporal organization, with older adults showing more spatially constrained and lateralized modulation patterns (Rayson et al., 2023). Beyond these burst characteristics, we found that age influenced the temporal organization of beta burst propagation globally across the cortex. With advancing age, burst onset occurred earlier during pre-movement but later during post-movement, creating an expanded temporal window of beta activity. This pattern was remarkably consistent across nearly all cortical regions, indicating that aging affects the timing of beta burst propagation broadly rather than in a region-specific manner. This temporal expansion may reflect age-related slowing of neural dynamics or altered cortical excitability that prolongs motor-related beta activity.

Additionally, this temporal expansion has functional implications. Mediation analysis revealed that age-related expansion of beta burst timing mediated the relationship between age and reaction time. The timing of beta dynamics directly impacts motor performance, with earlier desynchronization associated with faster reaction times (Toledo et al., 2016). The delayed post-movement burst offset in older adults may prolong motor inhibition, while earlier pre-movement onset may represent incomplete compensation. The observed expanded temporal window provides a mechanism linking cortical oscillatory timing to age-related behavioral slowing.

Several limitations should be noted. First, our analysis focused on cortical beta burst propagation and did not examine subcortical-to-cortical pathways, which are well-established routes for motor information flow (Diesburg et al., 2021). The optical flow technique, which analyzes propagation on a continuous surface, cannot readily integrate cortical and subcortical structures into a unified spatial framework due to their distinct anatomical geometries and spatial planes. Second, our trial-averaged approach was optimized for detecting spatial propagation patterns but may not capture trial-by-trial temporal variability that relates to behavioral performance (Little et al., 2019). Future studies may address both limitations by combining single-trial analyses with methods capable of integrating cortical and subcortical dynamics, providing a more comprehensive understanding of beta burst propagation across distributed motor networks and its relationship to behavior.

## Conclusion

We showed that beta burst activity propagates systematically along the posterior-anterior cortical axis during motor control. This propagation was task-specific, exhibited hemispheric asymmetry, and reversed direction between movement phases, reflecting distinct the functional demands of motor preparation versus motor state stabilization. Propagation patterns were associated with cortical neurochemical architecture in a hemisphere-specific and phase-dependent manner. Age-related temporal expansion of beta burst activity mediated the relationship between age and motor slowing. Overall, these findings highlight beta burst propagation as an organized feature of motor cortex dynamics, shaped by cortical architecture and modulated across the lifespan.

## References

Adam, R., Isabella, S., & Chan, J. L. (2015). Insight into motor control and motor impairment from stroke and beta oscillations. Journal of Neurophysiology, 114(6), 3033–3035. 10.1152/jn.00098.2015

Alahmadi, A. (2024). Functional Connectivity Profiles of Ten Sub-Regions within the Premotor and Supplementary Motor Areas: Insights into Neurophysiological Integration. Diagnostics, 14(17), 1990. https://www.mdpi.com/2075-4418/14/17/1990

Bezard, E., Li, Q., Hulme, H., Fridjonsdottir, E., Nilsson, A., Pioli, E., Andren, P. E., & Crossman, A. R. (2020). µ Opioid Receptor Agonism for L-DOPA-Induced Dyskinesia in Parkinson’s Disease. The Journal of Neuroscience, 40(35), 6812–6819. 10.1523/jneurosci.0610-20.2020

Brady, B., Power, L., & Bardouille, T. (2020). Age-related trends in neuromagnetic transient beta burst characteristics during a sensorimotor task and rest in the Cam-CAN open-access dataset. NeuroImage, 222, 117245. 10.1016/j.neuroimage.2020.117245

Brovelli, A., Ding, M., Ledberg, A., Chen, Y., Nakamura, R., & Bressler, S. L. (2004). Beta oscillations in a large-scale sensorimotor cortical network: Directional influences revealed by Granger causality. Proceedings of the National Academy of Sciences, 101(26), 9849–9854. doi:10.1073/pnas.0308538101

Cahalane, D. J., Charvet, C. J., & Finlay, B. L. (2012). Systematic, balancing gradients in neuron density and number across the primate isocortex. Frontiers in neuroanatomy, 6, 28.

Charvet, C. J., & Finlay, B. L. (2014). Evo-Devo and the Primate Isocortex: The Central Organizing Role of Intrinsic Gradients of Neurogenesis. Brain Behavior and Evolution, 84(2), 81–92. 10.1159/000365181

Chikermane, M., Weerdmeester, L., Rajamani, N., Köhler, R. M., Merk, T., Vanhoecke, J., Horn, A., & Neumann, W. J. (2024). Cortical beta oscillations map to shared brain networks modulated by dopamine. eLife, 13, RP97184. 10.7554/eLife.97184

Debaere, F., Wenderoth, N., Sunaert, S., Van Hecke, P., & Swinnen, S. P. (2003). Internal vs external generation of movements: differential neural pathways involved in bimanual coordination performed in the presence or absence of augmented visual feedback. NeuroImage, 19(3), 764–776. 10.1016/S1053-8119(03)00148-4

Diesburg, D. A., Greenlee, J. D. W., & Wessel, J. R. (2021). Cortico-subcortical β burst dynamics underlying movement cancellation in humans. eLife, 10, e70270. 10.7554/eLife.70270

Donoghue, T., Haller, M., Peterson, E. J., Varma, P., Sebastian, P., Gao, R., Noto, T., Lara, A. H., Wallis, J. D., Knight, R. T., Shestyuk, A., & Voytek, B. (2020). Parameterizing neural power spectra into periodic and aperiodic components. Nature Neuroscience, 23(12), 1655–1665. 10.1038/s41593-020-00744-x

Elston, G. N. (2000). Pyramidal cells of the frontal lobe: all the more spinous to think with. The Journal of Neuroscience, 20(18), RC95.

Engel, A. K., & Fries, P. (2010). Beta-band oscillations—signalling the status quo? Current Opinion in Neurobiology, 20(2), 156–165. 10.1016/j.conb.2010.02.015

Feingold, J., Gibson, D. J., DePasquale, B., & Graybiel, A. M. (2015). Bursts of beta oscillation differentiate postperformance activity in the striatum and motor cortex of monkeys performing movement tasks. Proceedings of the National Academy of Sciences, 112(44), 13687–13692. doi:10.1073/pnas.1517629112

Fischl, B. (2012). FreeSurfer. NeuroImage, 62(2), 774–781. 10.1016/j.neuroimage.2012.01.021

Fischl, B., Sereno, M. I., Tootell, R. B. H., & Dale, A. M. (1999). High-resolution intersubject averaging and a coordinate system for the cortical surface. Human Brain Mapping, 8(4), 272–284. 10.1002/(SICI)1097-0193(1999)8:4<272::AID-HBM10>3.0.CO;2-4

Fries, P. (2005). A mechanism for cognitive dynamics: neuronal communication through neuronal coherence. Trends in Cognitive Sciences, 9(10), 474–480. 10.1016/j.tics.2005.08.011

Gaetz, W., Edgar, J. C., Wang, D. J., & Roberts, T. P. L. (2011). Relating MEG measured motor cortical oscillations to resting γ-Aminobutyric acid (GABA) concentration. NeuroImage, 55(2), 616–621. 10.1016/j.neuroimage.2010.12.077

Gross, J., Pollok, B., Dirks, M., Timmermann, L., Butz, M., & Schnitzler, A. (2005). Task-dependent oscillations during unimanual and bimanual movements in the human primary motor cortex and SMA studied with magnetoencephalography. NeuroImage, 26(1), 91–98. 10.1016/j.neuroimage.2005.01.025

Hall, S. D., Stanford, I. M., Yamawaki, N., McAllister, C. J., Rönnqvist, K. C., Woodhall, G. L., & Furlong, P. L. (2011). The role of GABAergic modulation in motor function related neuronal network activity. NeuroImage, 56(3), 1506–1510. 10.1016/j.neuroimage.2011.02.025

Hansen, J. Y., Shafiei, G., Markello, R. D., Smart, K., Cox, S. M. L., Nørgaard, M., Beliveau, V., Wu, Y., Gallezot, J.-D., Aumont, É., Servaes, S., Scala, S. G., DuBois, J. M., Wainstein, G., Bezgin, G., Funck, T., Schmitz, T. W., Spreng, R. N., Galovic, M., … Misic, B. (2022). Mapping neurotransmitter systems to the structural and functional organization of the human neocortex. Nature Neuroscience, 25(11), 1569–1581. 10.1038/s41593-022-01186-3

Heinrichs-Graham, E., Kurz, M. J., Gehringer, J. E., & Wilson, T. W. (2017). The functional role of post-movement beta oscillations in motor termination. Brain Structure and Function, 222(7), 3075–3086. 10.1007/s00429-017-1387-1

Heinrichs-Graham, E., McDermott, T. J., Mills, M. S., Wiesman, A. I., Wang, Y.-P., Stephen, J. M., Calhoun, V. D., & Wilson, T. W. (2018). The lifespan trajectory of neural oscillatory activity in the motor system. Developmental Cognitive Neuroscience, 30, 159–168. 10.1016/j.dcn.2018.02.013

Hillebrand, A., Tewarie, P., van Dellen, E., Yu, M., Carbo, E. W. S., Douw, L., Gouw, A. A., van Straaten, E. C. W., & Stam, C. J. (2016). Direction of information flow in large-scale resting-state networks is frequency-dependent. Proceedings of the National Academy of Sciences, 113(14), 3867–3872. doi:10.1073/pnas.1515657113

Hosaka, R., Nakajima, T., Aihara, K., Yamaguchi, Y., & Mushiake, H. (2015). Arm-use dependent lateralization of gamma and beta oscillations in primate medial motor areas. Neural Networks, 62, 62–66. 10.1016/j.neunet.2014.06.004

Huntenburg, J. M., Bazin, P.-L., & Margulies, D. S. (2018). Large-Scale Gradients in Human Cortical Organization. Trends in Cognitive Sciences, 22(1), 21–31. 10.1016/j.tics.2017.11.002

Joviano-Santos, J. V., Kljakic, O., Magalhães-Gomes, M. P. S., Valadão, P. A. C., de Oliveira, L. R., Prado, M. A. M., Prado, V. F., & Guatimosim, C. (2021). Motoneuron-specific loss of VAChT mimics neuromuscular defects seen in congenital myasthenic syndrome. The FEBS Journal, 288(18), 5331–5349. 10.1111/febs.15825

Lefèvre, J., & Baillet, S. (2008). Optical Flow and Advection on 2-Riemannian Manifolds: A Common Framework. IEEE Transactions on Pattern Analysis and Machine Intelligence, 30(6), 1081–1092. 10.1109/TPAMI.2008.51

Lefèvre, J., & Baillet, S. (2009). Optical flow approaches to the identification of brain dynamics. Human Brain Mapping, 30(6), 1887–1897. 10.1002/hbm.20781

Lemon, R. N. (2008). Descending Pathways in Motor Control. Annual Review of Neuroscience, 31(Volume 31, 2008), 195-218. 10.1146/annurev.neuro.31.060407.125547

Little, S., Bonaiuto, J., Barnes, G., & Bestmann, S. (2019). Human motor cortical beta bursts relate to movement planning and response errors. PLOS Biology, 17(10), e3000479. 10.1371/journal.pbio.3000479

Liu, X., Wiesman, A. I., & Baillet, S. (2026). Hierarchical Flows of Human Cortical Activity. bioRxiv, 2026.2003.2019.712872. 10.64898/2026.03.19.712872

Lundqvist, M., Miller, E. K., Nordmark, J., Liljefors, J., & Herman, P. (2024). Beta: bursts of cognition. Trends in Cognitive Sciences, 28(7), 662–676. 10.1016/j.tics.2024.03.010

Mahjoory, K., Schoffelen, J.-M., Keitel, A., & Gross, J. (2020). The frequency gradient of human resting-state brain oscillations follows cortical hierarchies. eLife, 9, e53715. 10.7554/eLife.53715

Margulies, D. S., Ghosh, S. S., Goulas, A., Falkiewicz, M., Huntenburg, J. M., Langs, G., Bezgin, G., Eickhoff, S. B., Castellanos, F. X., Petrides, M., Jefferies, E., & Smallwood, J. (2016). Situating the default-mode network along a principal gradient of macroscale cortical organization. Proceedings of the National Academy of Sciences, 113(44), 12574–12579. doi:10.1073/pnas.1608282113

Markello, R. D., & Misic, B. (2021). Comparing spatial null models for brain maps. NeuroImage, 236, 118052. 10.1016/j.neuroimage.2021.118052

Nagao, M., Misawa, H., Kato, S., & Hirai, S. (1998). Loss of Cholinergic Synapses on the Spinal Motor Neurons of Amyotrophic Lateral Sclerosis. Journal of Neuropathology & Experimental Neurology, 57(4), 329–333. 10.1097/00005072-199804000-00004

Nowak, M., Hinson, E., van Ede, F., Pogosyan, A., Guerra, A., Quinn, A., Brown, P., & Stagg, C. J. (2017). Driving Human Motor Cortical Oscillations Leads to Behaviorally Relevant Changes in Local GABA A Inhibition: A tACS-TMS Study. The Journal of Neuroscience, 37(17), 4481–4492. 10.1523/jneurosci.0098-17.2017

Paquola, C., Vos De Wael, R., Wagstyl, K., Bethlehem, R. A. I., Hong, S.-J., Seidlitz, J., Bullmore, E. T., Evans, A. C., Misic, B., Margulies, D. S., Smallwood, J., & Bernhardt, B. C. (2019). Microstructural and functional gradients are increasingly dissociated in transmodal cortices. PLOS Biology, 17(5), e3000284. 10.1371/journal.pbio.3000284

Pfurtscheller, G., Graimann, B., Huggins, J. E., Levine, S. P., & Schuh, L. A. (2003). Spatiotemporal patterns of beta desynchronization and gamma synchronization in corticographic data during self-paced movement. Clinical Neurophysiology, 114(7), 1226–1236. 10.1016/S1388-2457(03)00067-1

Picazio, S., Veniero, D., Ponzo, V., Caltagirone, C., Gross, J., Thut, G., & Koch, G. (2014). Prefrontal Control over Motor Cortex Cycles at Beta Frequency during Movement Inhibition. Current Biology, 24(24), 2940–2945. 10.1016/j.cub.2014.10.043

Power, L., & Bardouille, T. (2021). Age-related trends in the cortical sources of transient beta bursts during a sensorimotor task and rest. NeuroImage, 245, 118670. 10.1016/j.neuroimage.2021.118670

Rayson, H., Szul, M. J., El-Khoueiry, P., Debnath, R., Gautier-Martins, M., Ferrari, P. F., Fox, N., & Bonaiuto, J. J. (2023). Bursting with Potential: How Sensorimotor Beta Bursts Develop from Infancy to Adulthood. The Journal of Neuroscience, 43(49), 8487–8503. 10.1523/jneurosci.0886-23.2023

Rossini, P. M., Rossi, S., Babiloni, C., & Polich, J. (2007). Clinical neurophysiology of aging brain: From normal aging to neurodegeneration. Progress in Neurobiology, 83(6), 375–400. 10.1016/j.pneurobio.2007.07.010

Rossiter, H. E., Davis, E. M., Clark, E. V., Boudrias, M.-H., & Ward, N. S. (2014). Beta oscillations reflect changes in motor cortex inhibition in healthy ageing. NeuroImage, 91, 360–365. 10.1016/j.neuroimage.2014.01.012

Rubino, D., Robbins, K. A., & Hatsopoulos, N. G. (2006). Propagating waves mediate information transfer in the motor cortex. Nature Neuroscience, 9(12), 1549–1557. 10.1038/nn1802

Schmidt, R., Herrojo Ruiz, M., Kilavik, B. E., Lundqvist, M., Starr, P. A., & Aron, A. R. (2019). Beta Oscillations in Working Memory, Executive Control of Movement and Thought, and Sensorimotor Function. The Journal of Neuroscience, 39(42), 8231–8238. 10.1523/jneurosci.1163-19.2019

Schütz, B. (2005). Imbalanced excitatory to inhibitory synaptic input precedes motor neuron degeneration in an animal model of amyotrophic lateral sclerosis. Neurobiology of Disease, 20(1), 131–140. 10.1016/j.nbd.2005.02.006

Shafto, M. A., Tyler, L. K., Dixon, M., Taylor, J. R., Rowe, J. B., Cusack, R., Calder, A. J., Marslen-Wilson, W. D., Duncan, J., Dalgleish, T., Henson, R. N., Brayne, C., Matthews, F. E., & Cam, C. A. N. (2014). The Cambridge Centre for Ageing and Neuroscience (Cam-CAN) study protocol: a cross-sectional, lifespan, multidisciplinary examination of healthy cognitive ageing. BMC Neurology, 14(1), 204. 10.1186/s12883-014-0204-1

Sherman, M. A., Lee, S., Law, R., Haegens, S., Thorn, C. A., Hämäläinen, M. S., Moore, C. I., & Jones, S. R. (2016). Neural mechanisms of transient neocortical beta rhythms: Converging evidence from humans, computational modeling, monkeys, and mice. Proceedings of the National Academy of Sciences, 113(33), E4885–E4894. doi:10.1073/pnas.1604135113

Shin, H., Law, R., Tsutsui, S., Moore, C. I., & Jones, S. R. (2017). The rate of transient beta frequency events predicts behavior across tasks and species. eLife, 6, e29086. 10.7554/eLife.29086

Szul, M. J., Papadopoulos, S., Alavizadeh, S., Daligaut, S., Schwartz, D., Mattout, J., & Bonaiuto, J. J. (2023). Diverse beta burst waveform motifs characterize movement-related cortical dynamics. Progress in Neurobiology, 228, 102490. 10.1016/j.pneurobio.2023.102490

Tadel, F., Baillet, S., Mosher, J. C., Pantazis, D., & Leahy, R. M. (2011). Brainstorm: a user-friendly application for MEG/EEG analysis. *Intell*. Neuroscience, 2011, Article 8. 10.1155/2011/879716

Taylor, J. R., Williams, N., Cusack, R., Auer, T., Shafto, M. A., Dixon, M., Tyler, L. K., Cam, C. A. N., & Henson, R. N. (2017). The Cambridge Centre for Ageing and Neuroscience (Cam-CAN) data repository: Structural and functional MRI, MEG, and cognitive data from a cross-sectional adult lifespan sample. NeuroImage, 144, 262–269. 10.1016/j.neuroimage.2015.09.018

Toledo, D. R., Manzano, G. M., Barela, J. A., & Kohn, A. F. (2016). Cortical correlates of response time slowing in older adults: ERP and ERD/ERS analyses during passive ankle movement. Clinical Neurophysiology, 127(1), 655–663. 10.1016/j.clinph.2015.05.003

Torrecillos, F., Tinkhauser, G., Fischer, P., Green, A. L., Aziz, T. Z., Foltynie, T., Limousin, P., Zrinzo, L., Ashkan, K., Brown, P., & Tan, H. (2018). Modulation of Beta Bursts in the Subthalamic Nucleus Predicts Motor Performance. The Journal of Neuroscience, 38(41), 8905–8917. 10.1523/jneurosci.1314-18.2018

van Wijk, B. C. M., Beek, P. J., & Daffertshofer, A. (2012). Differential modulations of ipsilateral and contralateral beta (de)synchronization during unimanual force production. European Journal of Neuroscience, 36(1), 2088–2097. 10.1111/j.1460-9568.2012.08122.x

Veen, B. D. V., Drongelen, W. V., Yuchtman, M., & Suzuki, A. (1997). Localization of brain electrical activity via linearly constrained minimum variance spatial filtering. IEEE Transactions on Biomedical Engineering, 44(9), 867–880. 10.1109/10.623056

Wagner, J., Wessel, J. R., Ghahremani, A., & Aron, A. R. (2018). Establishing a Right Frontal Beta Signature for Stopping Action in Scalp EEG: Implications for Testing Inhibitory Control in Other Task Contexts. Journal of Cognitive Neuroscience, 30(1), 107–118. 10.1162/jocn_a_01183

Wessel, J. R. (2020). β-Bursts Reveal the Trial-to-Trial Dynamics of Movement Initiation and Cancellation. The Journal of Neuroscience, 40(2), 411–423. 10.1523/jneurosci.1887-19.2019

West, T. O., Duchet, B., Farmer, S. F., Friston, K. J., & Cagnan, H. (2023). When do bursts matter in the primary motor cortex? Investigating changes in the intermittencies of beta rhythms associated with movement states. Progress in Neurobiology, 221, 102397. 10.1016/j.pneurobio.2022.102397

Wilson, L. E., da Silva Castanheira, J., & Baillet, S. (2022). Time-resolved parameterization of aperiodic and periodic brain activity. eLife, 11, e77348. 10.7554/eLife.77348

Wiskerke, J., Van Mourik, Y., Schetters, D., Schoffelmeer, A. N., & Pattij, T. (2012). On the Role of Cannabinoid CB1- and μ-Opioid Receptors in Motor Impulsivity [Original Research]. Frontiers in Pharmacology, 3. 10.3389/fphar.2012.00108

Xifra-Porxas, A., Niso, G., Larivière, S., Kassinopoulos, M., Baillet, S., Mitsis, G. D., & Boudrias, M.-H. (2019). Older adults exhibit a more pronounced modulation of beta oscillations when performing sustained and dynamic handgrips. NeuroImage, 201, 116037. 10.1016/j.neuroimage.2019.116037

Zhang, X., Wang, H., Guo, Y., & Long, J. (2024). Beta rebound reduces subsequent movement preparation time by modulating of GABAA inhibition. Cerebral Cortex, 34(2). 10.1093/cercor/bhae037

Zhao, X., Lynch, J. G., Jr., & Chen, Q. (2010). Reconsidering Baron and Kenny: Myths and Truths about Mediation Analysis. Journal of Consumer Research, 37(2), 197–206. 10.1086/651257

Zich, C., Quinn, A. J., Bonaiuto, J. J., O’Neill, G., Mardell, L. C., Ward, N. S., & Bestmann, S. (2023). Spatiotemporal organisation of human sensorimotor beta burst activity. eLife, 12, e80160. 10.7554/eLife.80160

