## Supplementary materials for "Spatiotemporal Propagation of Sensorimotor Beta Bursts Across Adulthood"

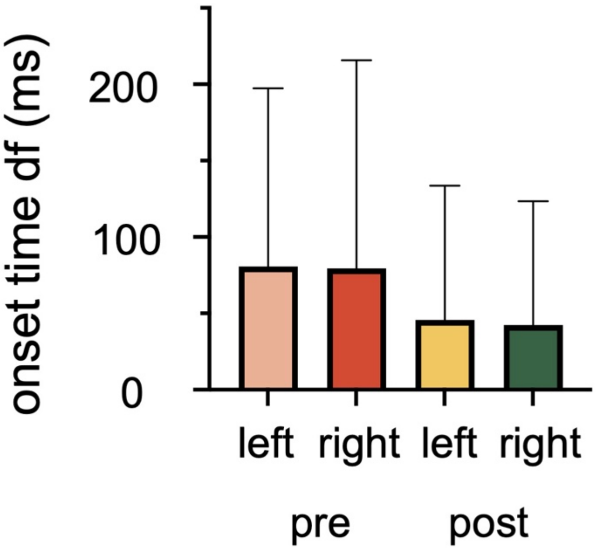


**Fig. S1.** Temporal spread of regional beta burst onset times. Bars show the difference between latest and earliest onset times (max - min) across cortical regions for each hemisphere and phase. Error bars: SD across participants.

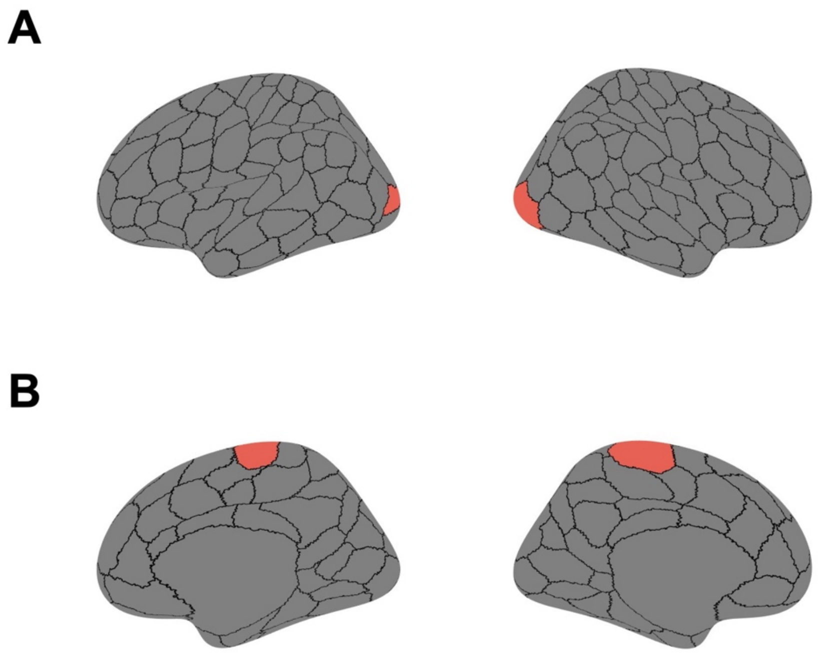

**Fig. S2.** Reference regions for computing anatomical distances. **(A)** Visual cortex reference region (red, lateral views) used to compute posterior-anterior distances. **(B)** Motor cortex reference region (red, medial views) used to compute central-peripheral distances.

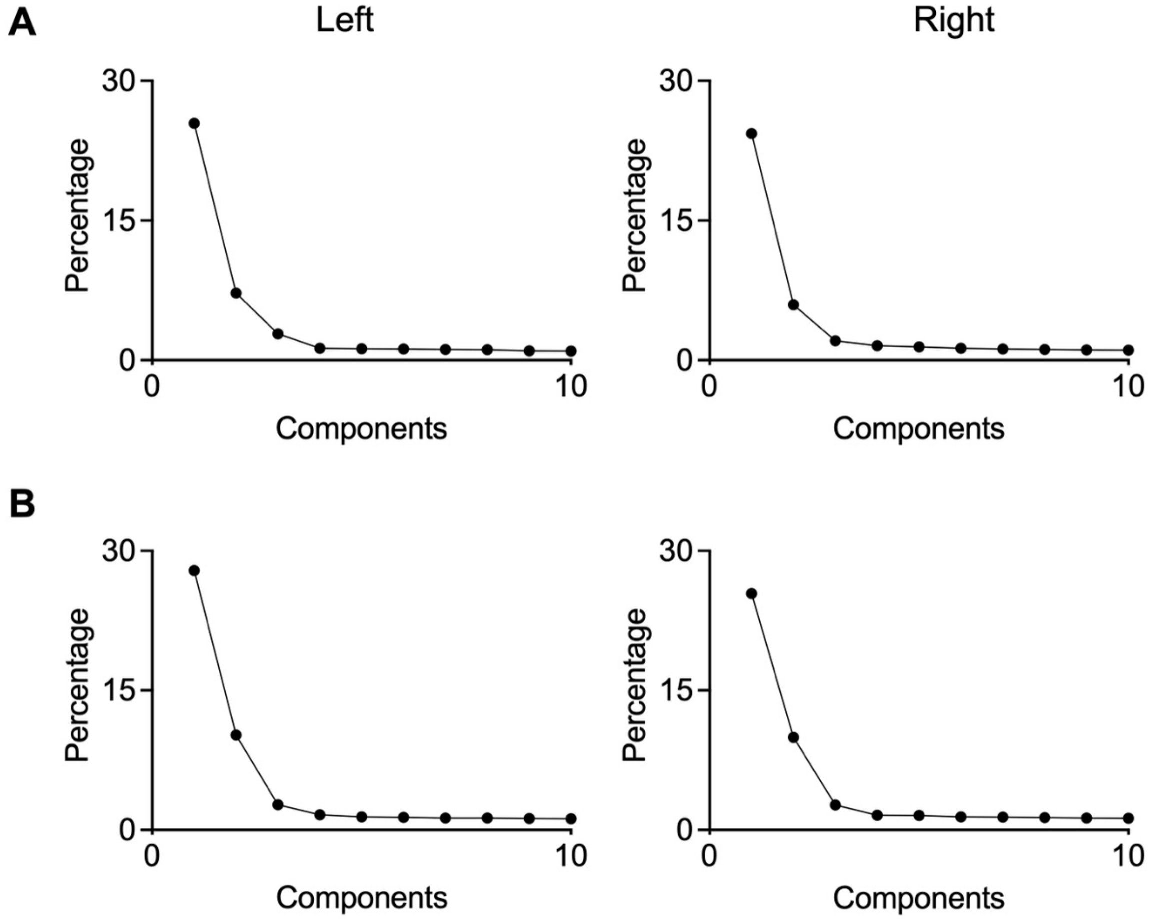


**Fig. S3.** Variance explained by principal components of beta burst onset timing. Percentage of variance explained by the first 10 principal components derived from PCA of beta burst onset time ranks. **(A)** Pre-movement period. **(B)** Post-movement period.

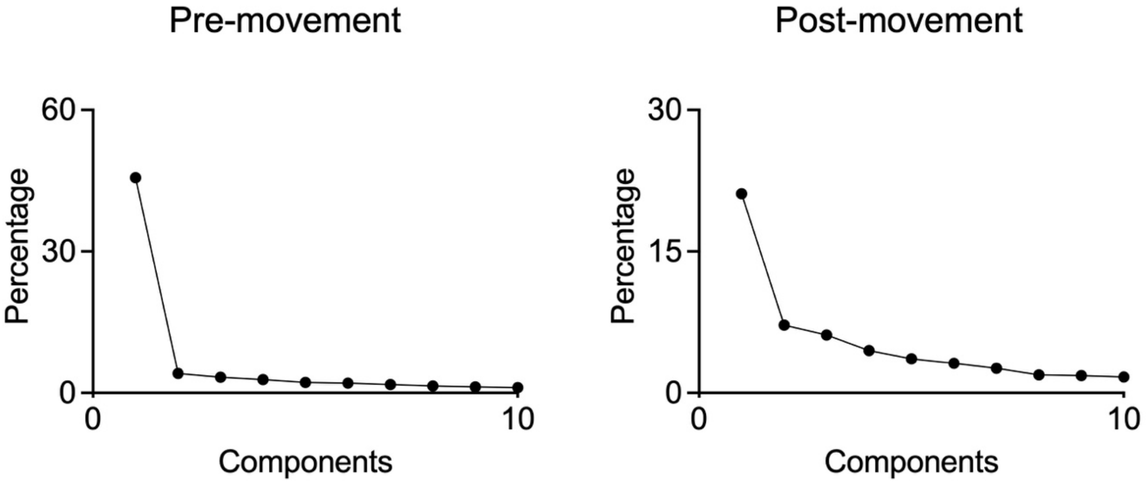


**Fig. S4.** Percentage of variance explained by the first 10 principal components derived from PCA of optical flow energy ranks.

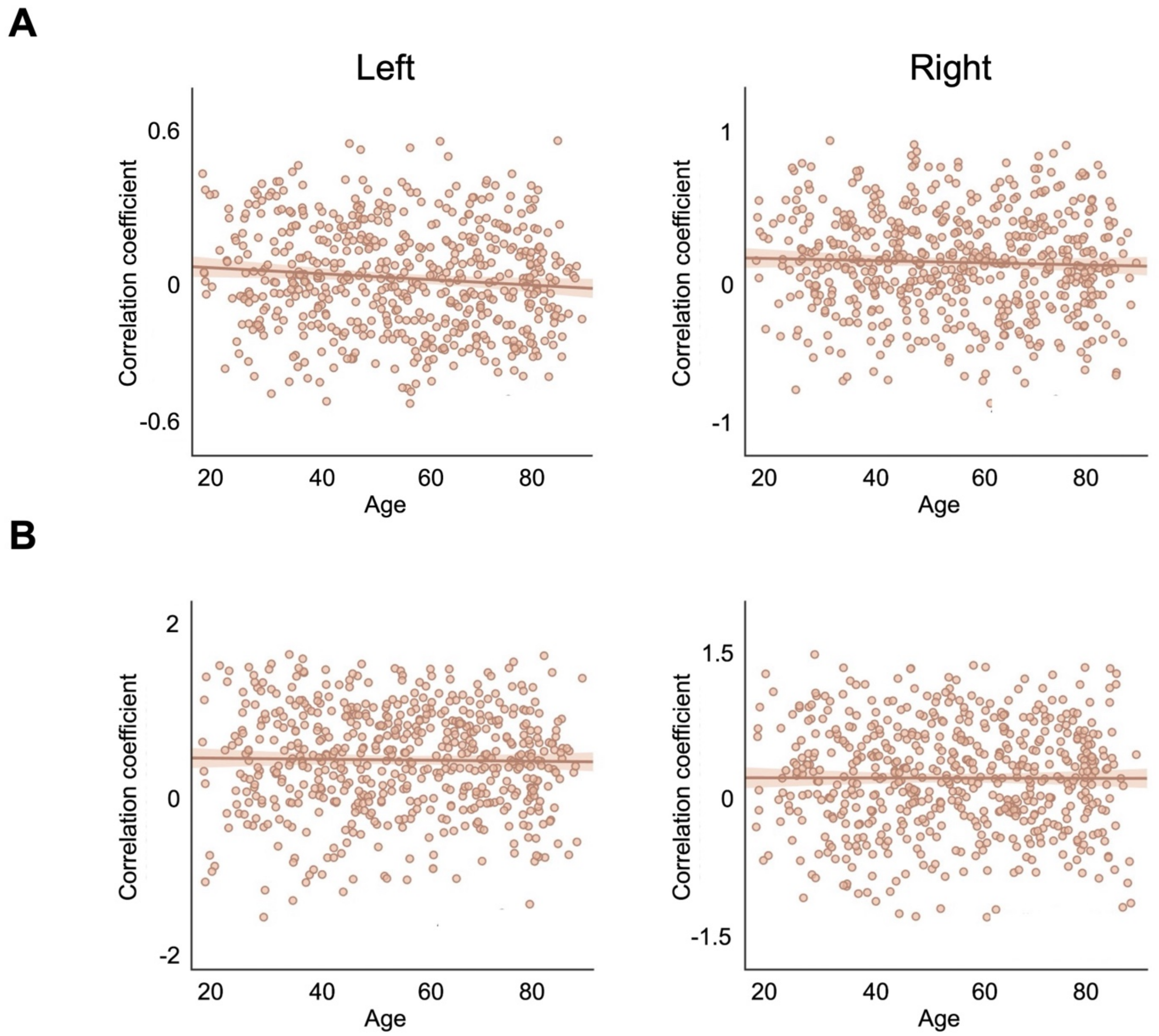


**Fig. S5.** Correlation between age and beta burst onset time-PC1 relationship across hemispheres and task phases. Correlation coefficients (Fisher z-transformed) between individual participants' beta burst onset time rank and group-level PC1 plotted against age for left (left panel) and right (right panel) hemispheres. **(A)** Pre-movement phase. **(B)** Post-movement phase. No significant correlation was observed among all plots.
